# The sodium leak channel NALCN encodes the major background sodium ion conductance in mouse anterior pituitary cells

**DOI:** 10.1101/2021.08.02.454810

**Authors:** Marziyeh Belal, Mariusz Mucha, Arnaud Monteil, Paul G Winyard, Robert Pawlak, Jamie J. Walker, Joel Tabak, Mino D C Belle

**Author notes:** **Author Contributions:** M.B, M.M, A.M, P.G.W, R.P, J.J.W, J.T and M.D.C.B conceived and designed research; M.B and M.M performed genetic manipulation experiments and M.B performed electrophysiology experiments as well as calcium imaging and data analysis; M.B, A.M, J.J.W, J.T and M.D.C.B wrote the paper. **Competing Interest Statement:** The authors declare no competing interests.

## Abstract

The pituitary gland produces and secretes a variety of hormones that are essential to life, such as for the regulation of growth and development, metabolism, reproduction, and the stress response. This is achieved through an intricate signalling interplay between the brain and peripheral feedback signals that shapes pituitary cell excitability by regulating ion channel properties of these cells. In addition, endocrine anterior pituitary cells fire action potentials spontaneously to regulate intracellular calcium ([Ca^2+^]_i_) level, an essential signalling conduit for hormonal secretion. To this end, pituitary cells have to critically regulate their resting membrane potential (RMP) close to firing threshold, but the molecular identity of the ionic mechanisms involved remains largely unknown. Here, we revealed that the sodium leak channel NALCN, known to modulate neuronal excitability elsewhere in the brain, acts to regulate excitability in the mouse anterior endocrine pituitary cells. Using viral transduction combined with powerful electrophysiology methods and calcium imaging, we show that NALCN forms the major Na^+^ leak conductance in these cells, appropriately tuning cellular RMP for sustaining spontaneous firing activity. Genetic interruption of NALCN channel activity drastically hyperpolarised the cells, suppressing firing and ([Ca^2+^]_i_) oscillations. Remarkably, we uncover that NALCN conductance formed a very small fraction of the total cell conductance, but yet had a profound impact on pituitary cell excitability. Our results also provide a possible mechanism through which hypothalamic and hormone feedback signals can powerfully affect pituitary activity to influence hormonal function.

## Introduction

The electrical activity of excitable cells appropriately responds to signals that are externally driven and/or those that are intrinsically derived to support essential physiological functions, such as breathing, heartbeat, and hormone release (Cui et al, 2016; Protze et al, 2017; Rorsman and Ashcroft, 2018; Bertram et al, 2018). Such excitability capacity relies on depolarising conductances that appropriately maintain the resting membrane potential (RMP) of cells near their firing threshold to support the spontaneous discharge of action potentials (APs). In hormone-secreting cells of the anterior pituitary gland, spontaneous AP firing results in rhythmic Ca^2+^ entry through voltage-gated calcium channels. These cytosolic/intracellular calcium ([Ca^2+^]_i_) oscillations serve a plethora of key physiological purposes, such as maintaining Ca^2+^ levels in intracellular calcium stores, regulating gene expression as well as evoking hormonal secretion (Mollard & Schlegel, 1996; Kwiecien & Hammond, 1998; Stojilkovic, 2012). Indeed, the silencing of spontaneous firing in endocrine pituitary cells abolishes [Ca^2+^]_i_ oscillations and hormone secretion (Kucka et al, 2010).

The ability for pituitary cells to produce spontaneous APs is in part due to the maintenance of their depolarised RMP relative to the much-hyperpolarised K^+^ equilibrium potential (Fletcher et al, 2018). Replacing extracellular Na^+^ with large impermeable cations, such as NMDG^+^, starves the cells to Na^+^ entry, suppressing the cell’s ability to depolarise. Indeed, such treatments hyperpolarises the RMP, abolishing firing activity and [Ca^2+^]_i_ oscillations (Simasko, 1994; Sankaranarayan & Simasko, 1996; Kwiecien and Hammond, 1998; Tsaneva-Atanasova et al, 2007; Kucka et al, 2010, 2012; Tomic et al, 2011; Liang et al, 2011; Zemkova et al, 2016; Kayano et al, 2019). This indicates that constitutively active inward-depolarising Na^+^-dependent currents/conductance in pituitary cells critically operates to maintenance of the RMP close to the firing threshold. Pharmacological and electrophysiological investigation of this inward leak current suggested it to be mediated by a TTX-insensitive, voltage-independent, and constitutively active Na^+^-permeable channel/conductance (Fletcher et al, 2018). However, the molecular identity of the channels responsible for such resting Na^+^ conductance in pituitary cells remains unknown.

The voltage-independent nonselective Na^+^ leak channel NALCN has emerged as the major background Na^+^-dependent conductance in several neuronal populations. For example, NALCN is essential for maintaining spontaneous AP firing in hippocampal neurons (Lu et al, 2007), GABAergic and dopaminergic neurons of the midbrain (Lutas et al, 2016; Philippart and Khaliq, 2018) and neurons of the suprachiasmatic nucleus (Flourakis et al, 2015). In the ventral respiratory neurons of the brain stem, NALCN activity facilitates rhythmic and carbon dioxide-stimulated breathing, as well as responsiveness to neuropeptides (Lu et al, 2007; Shi et al, 2016; Yeh et al, 2017), and NALCN regulate excitability in endocrine cells, such as pancreatic β cells (Swayne et al, 2009).

In the pituitary, a high level of NALCN mRNA expression is reported (Swayne et al, 2009); recent transcriptomic data also indicate that anterior pituitary cells express the NALCN gene and its known regulatory subunits (UNC-79, UNC-80, FAM155A) at level that is significantly higher when compared with gene expression of other known cationic leak channels, such as the TRP and HCN channels (Paul Le Tissier, Jacques Drouin and Patrice Mollard, personal communication, 04/2021). In addition, the pharmacological profile of NALCN in neurons is similar to that of resting Na^+^ conductance measured in pituitary cells: TTX-insensitive, extracellular Ca^2+^- and NMDG^+^-sensitive (Fletcher et al, 2018; Lu et al, 2007), and recent work has indicated that NALCN activity influences excitability in GH3 pituitary clonal cell line (Impheng et al, 2021).

Together, this strongly suggests that NALCN is the primary contributor to the background Na^+^ conductance and [Ca^2+^]_i_ oscillations in anterior endocrine pituitary cells, and consequently plays a key role in regulating pituitary cell excitability. To evaluate this, we used a parallel lentiviral-mediated NALCN knockdown strategy combined with powerful electrophysiology methods and calcium imaging in mouse primary anterior pituitary cells. Our results unequivocally revealed that NALCN is the main contributor to the background Na^+^-dependant depolarising conductance in pituitary cells, which action is key to maintain cellular excitability. Remarkably, we were also able to estimate the magnitude of this NALCN conductance.

## Results

### NALCN channel protein is expressed in primary mouse anterior pituitary cells

Previous research has revealed the expression of the *Nalcn* gene in mouse anterior pituitary gland at the mRNA level (Swayne et al, 2009). Thus, we first confirmed NALCN protein expression in mouse primary endocrine pituitary cells with a NALCN antibody directed against the extracellular epitope of the NALCN channel protein (Alomone Labs, Israel, #ASC-022). Our results revealed the presence of NALCN channel in mouse endocrine anterior pituitary cells (Figure 1), with the majority of the anterior pituitary cells stained for NALCN. No staining was observed when the NALCN antibody was omitted (Figure 1H).

**Figure 1.**
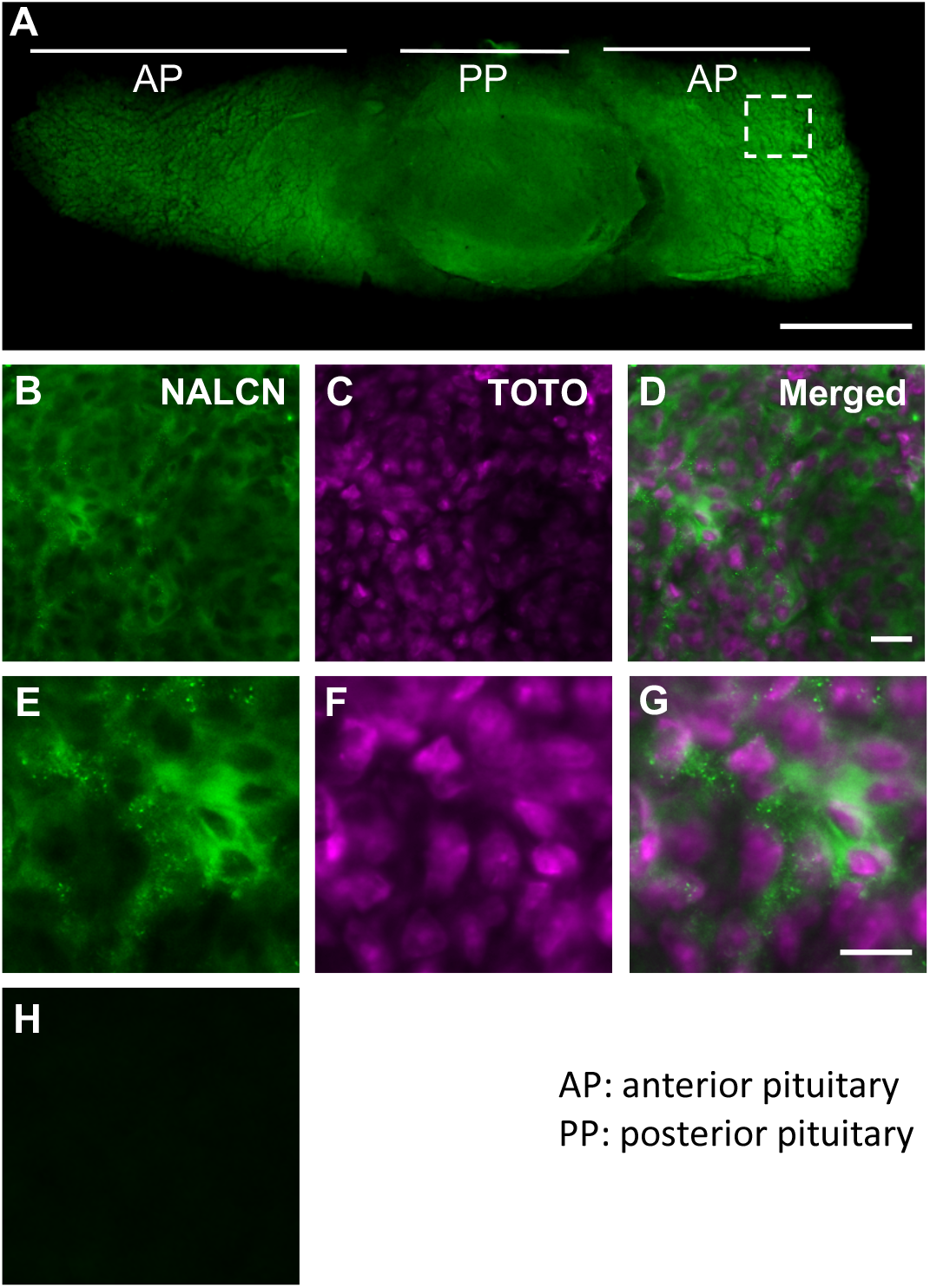
The mouse pituitary gland expresses NALCN ion channel. Immunofluorescence staining revealed the presence of NALCN channel protein in the mouse anterior pituitary gland (n=3). NALCN channel is shown in green. TOTO-3 (shown in magenta) was used to identify cell bodies by visualising cellular nucleic acids. **A)** Transverse section of a pituitary gland under 10X magnification excited with 488 nm light to detect NALCN channel immunofluorescence. **B)** NALCN channel visualisation in the area defined by dashed square in A. **C)** TOTO-labelled nucleic acid in the same area shown in B. **D)** Merged image from B and C. **B**, **C** and **D** are under 20X magnification. **E-G)** are under higher digital magnification using confocal microscopy. **H)** Negative control tissue for which the primary anti-NALCN antibody was omitted showing a lack of staining. Scale bars indicate 500 μm in panel A, and 20 μm in panels D and G.

### NALCN regulates spontaneous firing in primary pituitary cells

Although gadolinium (Gd^3+^) and flufenamic acid (FFA) act to suppress spontaneous firing in pituitary cells by causing hyperpolarisation (Kucka et al, 2012), and thus revealing the presence of a functional non-selective cationic conductance (NSCC) in these cells, it is noteworthy that Gd^3+^ and FFA are not specific blockers for a particular non-selective cationic conductance. Thus, NSCCs other than NALCN may contribute to the effects of FFA and Gd^3+^ on membrane potential and firing activity. To determine if NALCN is the main NSCC in endocrine anterior pituitary cells, we used a genetic manipulation approach to directly and selectively alter the activity of NALCN, investigating its role in the regulation of membrane potential and spontaneous firing in these cells. To this end, we used a lentiviral-mediated knockdown strategy to decrease NALCN channel expression/activity level in mouse endocrine pituitary cells (see methods), and then evaluated the resulting changes in electrophysiological properties of these cells.

Our data revealed that most, if not all, untreated control (untreated Ctrl) and scramble control (SCR Ctrl) cells exhibited spontaneous firing activity (Figure 2A, B, D, F), consistent with previous reports (*reviewed in* Fletcher et al, 2018). In contrast, 90% (28 of 31) of NALCN knockdown (NALCN KD) cells were found to be silent (Figure 2C, D, F), compared to just 17% (5 of 30) in SCR control and 19% (6 of 31) in untreated control cells (Figure 2D, F). In addition, NALCN KD cells exhibited a significantly more hyperpolarised resting membrane potential (RMP) compared both to untreated and SCR Ctrl cells (NALCN KD RMP: −63.6 ± 8.5 mV (mean ± SD), n=31, 12 animals; SCR Ctrl: - 45.2 mV ± 5.8 mV, n=30, 12 animals; untreated Ctrl: −47.5 mV ± 7.8, n=31, 12 animals; p<0.001, one-way ANOVA with post hoc Bonferroni correction; Figure 2E). There was no significant difference between the SCR Ctrl and untreated Ctrl cells, for neither the firing frequency (untreated Ctrl: 0.7 ± 0.6 Hz, n=25; SCR Ctrl: 0.5 ± 0.4 Hz, n=25, p>0.1, *t*-test; Figure 2F) nor the RMP (untreated Ctrl: −47.6 ± 7.6 mV, n=31; SCR Ctrl: −45.2 mV ± 5.8 mV, n=30 p>0.1, *t*-test; Figure 2E). Our results, therefore, indicate that NALCN activity in pituitary cells is critical for maintaining firing activity and sustaining of cellular RMP. In addition, since our results strongly indicated that the viral transduction *per se* did not affect the firing activity or RMP of pituitary cells, we henceforth made comparisons between NALCN KD and SCR Ctrl cells only.

**Figure 2.**
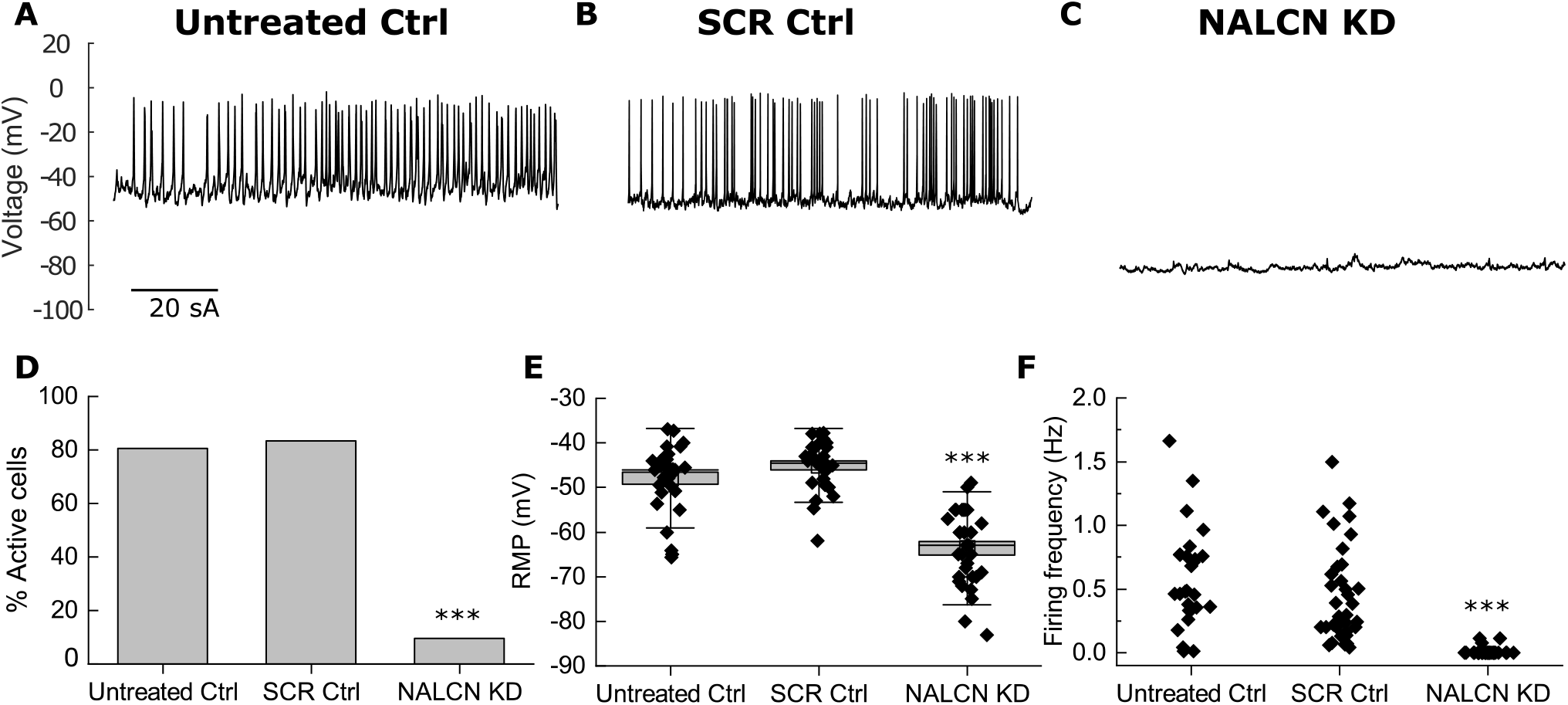
Knock-down of NALCN silences electrical activity in mouse pituitary cells. Representative traces of spontaneous firing activity from **A)** an untreated and **B)** a scramble control (SCR Ctrl) pituitary cell. **C)** Representative trace of electrical activity in a NALCN knockdown (NALCN KD) pituitary cell. **D)** Percentages of active and silent cells in untreated control, SCR Ctrl, and NALCN KD pituitary cells. **E)** Resting membrane potential of untreated control, SCR Ctrl, and NALCN KD pituitary cells (data presented as mean ± SD, untreated control: −47.6 ± 7.6 mV, n=31 vs SCR Ctrl: −45 mV ± 5.8 mV, n=30, p>0.1; NALCN KD: −63.6 ± 8.5 mV, n=31 vs SCR Ctrl and untreated control, p < 0.001, one-way ANOVA with Bonferroni correction). **F)** Distribution of firing frequency between untreated Ctrl, SCR Ctrl, and NALCN KD cells over a course of 600 seconds (data presented as mean ± SD, NALCN KD: 0 Hz, n= 31 vs SCR Ctrl: 0.5 ± 0.4 Hz, n= 25 and vs untreated Ctrl: 0.7 ± 0.6 Hz, n= 25, p < 0.001; SCR Ctrl vs untreated Ctrl: p > 0.1, one-way ANOVA with Bonferroni correction). Only 3/31 NALCN KD cells fired action potential, albeit at a lower frequency (0.1 ± 0.02 Hz) relative to the untreated and SCR controls. *** represents p < 0.001. In **(E)** boxes and whiskers represent the standard error of the mean and standard deviation, respectively. Each black dotes in **(E)** and **(F)** shows values from individual cells.

### Small conductance injection restores firing in NALCN KD primary pituitary cells

Having rendered pituitary cells hyperpolarised-silent with NALCN KD, we next used dynamic-clamp to examine whether the injection of nonselective cationic conductance to these silent NALCN KD cells could restore RMP and spontaneous firing activity. Indeed, hyperpolarised-silent NALCN KD cells became depolarised and restored typical firing activity and frequency when the non-selective cationic conductance was injected by very small amounts, such as 0.02 nS (Figure 3A). The added conductance required to bring NALCN KD cells to the typical RMP and firing activity/frequency observed in SCR Ctrl cells ranged from 0.02 to 0.12 nS (median 0.05 nS, n=28, Figure 3B).

**Figure 3.**
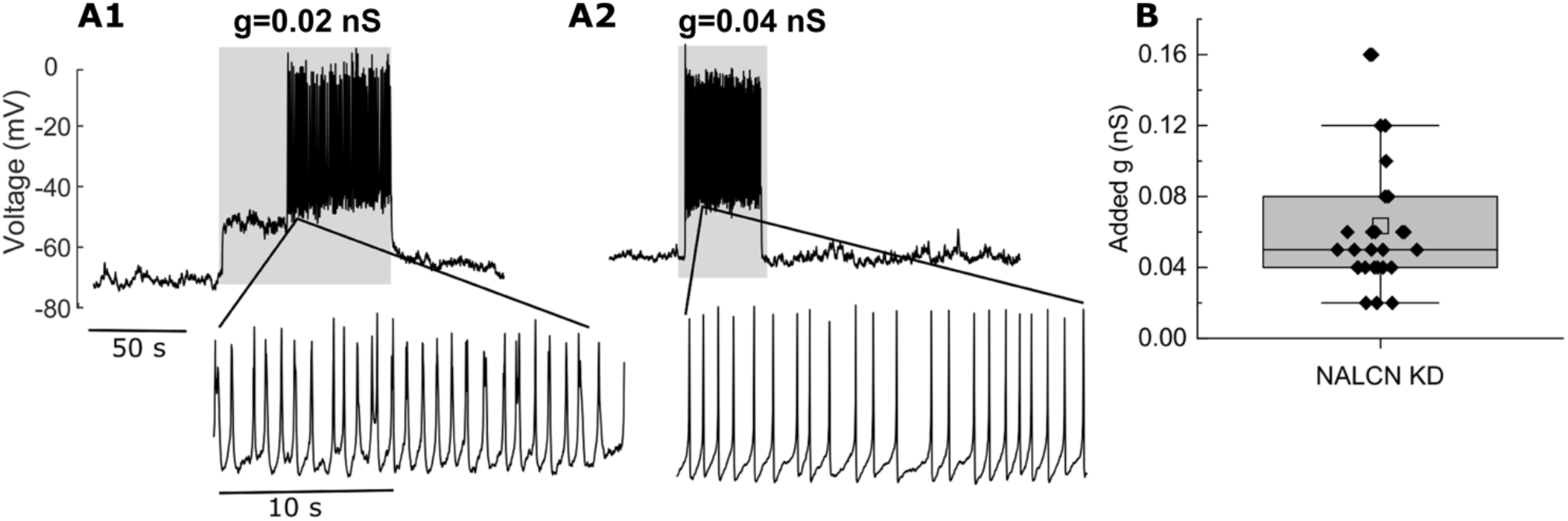
Dynamic clamp addition of nonselective cationic conductance restores firing activity in silent NALCN KD pituitary cells. **A)** Two representative voltage traces of silent NALCN KD pituitary cells. Firing activity was restored once the nonselective cationic conductance (g) was increased by amount as minute as 0.02 nS (e.g. silent cells depolarised and discharged barrages of action potentials when 0.02 or 0.04 nS conductance was applied (grey box), immediately returning to baseline after termination of added conductance. **B)** The distribution of added conductance (g) values required for restoring firing activity in hyperpolarised-silent NALCN KD pituitary cells. Median: 0.05 nS, n=28 from 12 animals.

Taken together, this first set of results indicate that NALCN is a key player in pituitary cell excitability, by modulating the RMP and subsequently, firing activity in primary pituitary cells. The amount of NALCN conductance lost by the cells after NALCN KD, as determined by dynamic clamp, was estimated to be on the order of 0.05 nS.

### NALCN majorly contributes to the inward Na^+^ leak currents in primary pituitary cells

To further determine the contribution of the NALCN-mediated inward leak current over other background Na^+^ currents in pituitary cells, we compared the inward current density in SCR Ctrl and NALCN KD cells while maintaining them at a holding potential of −80 mV, which in our setup also minimises any influence of K^+^ channel conductance. At −80 mV, the holding current density was significantly larger in the SCR Ctrl group compared to NALCN KD group (Figure 4A-C, SCR Ctrl (mean ± SD): −0.72 ± 0.2 pA/pF, n=15, 8 animals; NALCN KD: −0.19 ± 0.13 pA/pF, n=16, 8 animals; p< 0.001, *t*-test), confirming that the NALCN KD cells were indeed more hyperpolarised than their SCR Ctrl counterparts. Remarkably, this background Na^+^ conductance was reduced by 0.045 nS in SCR Ctrl, but by only 0.015 nS in NALCN KD (3 times smaller than what is measured in SCR Ctrl) when extracellular Na^+^ was substituted by the Na^+^ channel impermeant cation NMDG^+^. This indicates that the measured inward currents in SCR Ctrl were being mediated by a background Na^+^ conductance that was nearly absent in NALCN KD cells (Figure 4D, SCR Ctrl (median ± interquartile): 0.045 ± 0.02 nS, n = 16; vs NALCN KD: 0.015 ± 0.01 nS, n = 15; 8 animals in each condition, p < 0.001, Mann Whitney test).

**Figure 4.**
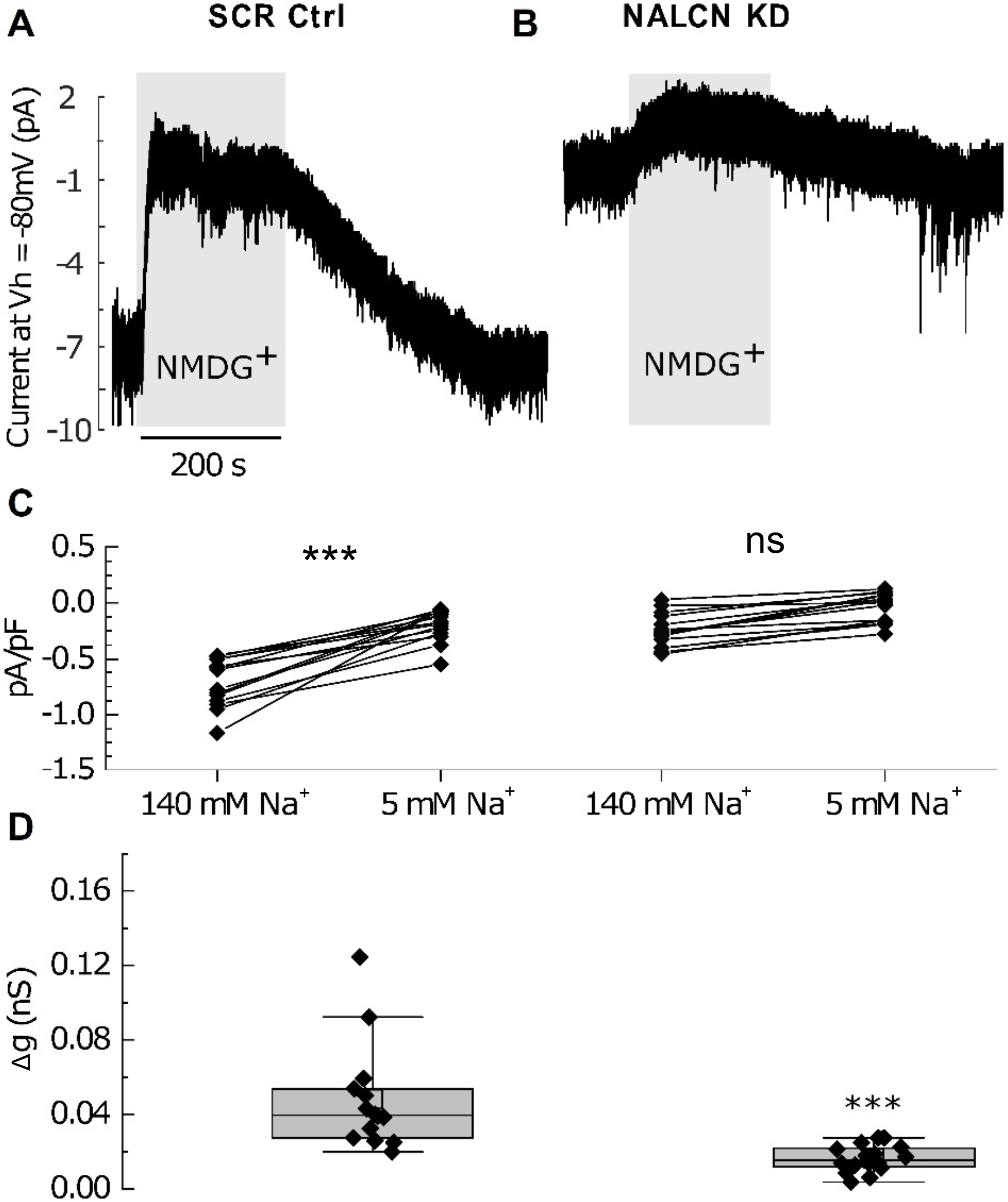
NALCN contributes to most of the inward Na^+^ leak conductance in endocrine anterior pituitary cells. **A-B)** Representative traces of inward Na^+^ leak currents (V_h_ = −80 mV) in a SCR Ctrl and NALCN KD pituitary cells as revealed by the substitution of extracellular Na^+^ with NMDG^+^ (grey boxes). **C)** The density of the inward Na^+^ leak current in NALCN KD cells was significantly reduced compared to SCR Ctrl (data presented as Δmean ± SD, SCR Ctrl: −0.72 ± 0.2 pA/pF (−5.2 ± 1.9 pA), n=15; NALCN KD: −0.19 ± 0.13 pA/pF (−1.5 ± 0.9 pA), n=16; 8 animals per condition, p< 0.001, *t*-test). **D)** Extracellular Na^+^ substitution with NMDG^+^ caused significant loss of background Na^+^ conductance in control but not in NALCN KD cells (data represented as ΔMedian ± interquartile, SCR control: 0.045 ± 0.02 nS, n = 16; NALCN KD: 0.015 ± 0.01 nS, n = 15; p < 0.001, Mann Whitney test).

These findings indicate that NALCN indeed contributes to most of the inward leak conductance in pituitary cells. Our results also provide a rough estimate of NALCN conductance magnitude in pituitary cells, by noting that the difference in background Na^+^ conductance between SCR Ctrl and NALCN KD cells is 0.03 nS (0.045 nS – 0.015 nS). This is probably an underestimate since some residual NALCN conductance may still be available in NALCN KD cells following the lentiviral knockdown. Importantly, our results identify that a change in background Na^+^ conductance of near 0.03 nS was enough to profoundly silence the electrical activity of endocrine anterior pituitary cell. This finding is consistent with endocrine pituitary cells having very large input resistance (~5 GΩ; Oxford and Dubinsky, 1988), meaning, very small variation in the current results in drastic changes in the RMP. This also agrees with our dynamic clamp observation where a 0.05 nS inward leak conductance injection was sufficient at fully restoring RMP and firing activity in NALCN KD cells (see above section and compare Figures 3 with 4).

### NALCN is required for spontaneous intracellular Ca^2+^ oscillations in primary pituitary cells

Previous studies have shown that the pattern of spontaneous firing determines the amplitude and duration of [Ca^2+^]_i_ oscillations in anterior endocrine pituitary cells (Stojilkovic et al, 2005, Stojilkovic et al, 2012), a necessary signalling link with excitability for hormonal secretion. Thus, we next tested whether NALCN KD affects spontaneous [Ca^2+^]_i_, oscillations in these cells. We found that approximately 57% (51/90) of SCR Ctrl pituitary cells exhibited spontaneous [Ca^2+^]_i_, oscillations (Figure 5A). In contrast, only 11% (4/36) of NALCN KD cells displayed such [Ca^2+^]_i_, transients (Figure 5), while the remaining 89% (32/36) were quiescent and did not generate any [Ca^2+^]_i_, oscillations (Figure 5B). The percentage (57%) of pituitary cells that produced [Ca^2+^]_i_, oscillations in Ctrl conditions was consistent with previous reports (Tomić et al, 2011). To quantitatively compare the size of the [Ca^2+^]_i_, oscillations between the SCR Ctrl and NALCN KD cells, the standard deviation (SD) of the baseline Ca^2+^ ratio trace for each individual cell was calculated over 600 seconds. Here, we use the size of the SD as a measure of the [Ca^2+^]_i_, oscillations/fluctuations magnitude (for example compare Figure 5A and B). Comparing the SD between the two groups revealed a significant difference between SCR Ctrl and NALCN KD cells (Figure 5C, SCR Ctrl: median=0.04, n=90; NALCN KD: median=0.015, n=36; p<0.001, Mann Whitney test, 5 animals per condition with each animal providing roughly equal number of cells), with [Ca^2+^]_i_, ratios showing larger fluctuations in SCR Ctrl cells when compared with NALCN KD cells. Together, this indicated that NALCN plays a key role in regulating [Ca^2+^]_i_, oscillations in primary pituitary cells. As in SCR Ctrl cells, challenging NALCN KD cells with high extracellular K^+^ (15 mM) induced depolarisation and consequently a rise in [Ca^2+^]_i_, ratios, serving as a positive control for these cells.

**Figure 5.**
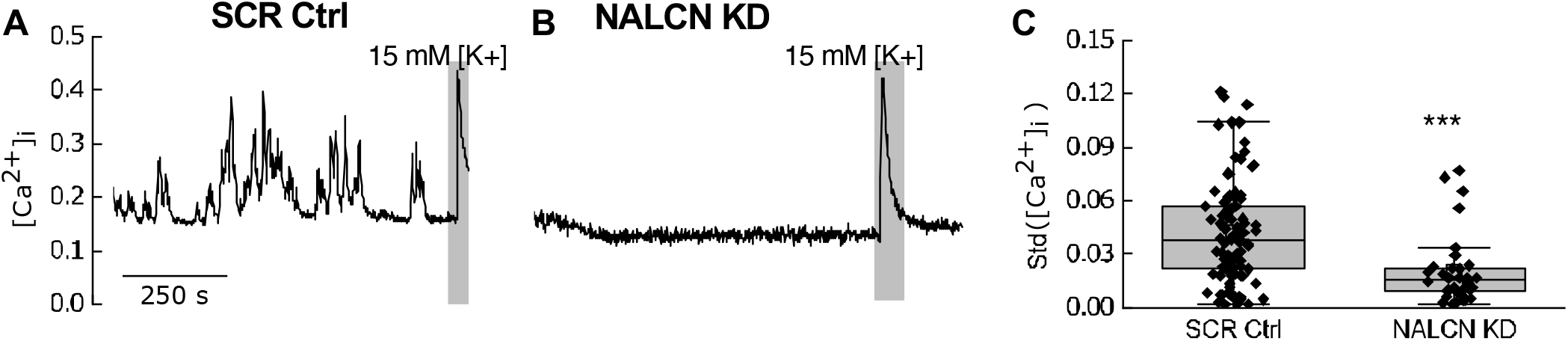
NALCN KD impacts intracellular Ca^2+^ transients. **A)** Representative trace of intracellular Ca^2+^ ([Ca^2+^]_i_) oscillations in an SCR control (SCR Ctrl) pituitary cell. **B)** A representative trace of [Ca^2+^]_i_ in a NALCN KD pituitary cell, showing abolished [Ca^2+^]_i_ oscillations. **C)** The standard deviation (Std) of each [Ca^2+^]_i_ ratio trace was calculated and statistical analysis revealed a significant alteration of [Ca^2+^]_i_ oscillations between the two groups (SCR Ctrl: median= 0.04, n = 90; NALCN KD: median= 0.015, n = 36, 5 mice per condition, p < 0.001, Mann Whitney test). Box: inter-quartile, Whiskers: range, excluding outliers, Central line: median. The last responses in panels A and B (grey boxes) represent [Ca^2+^]_i_ response to a rise in extracellular K^+^ (15 mM), used here as positive control to report cell viability.

### NALCN mediates a low extracellular Ca^2+^-induced depolarisation in primary pituitary cells

Removal of extracellular Ca^2+^ ([Ca^2+^]_e_) activates a low and sustained excitatory inward leak current in hippocampal neurons (Chu et al, 2003), by a mechanism that directly involves NALCN, acting through the UNC80-NALCN complex to regulate excitability (Lu et al, 2010). It is reported that in primary anterior pituitary cells, lowering or elimination of [Ca^2+^]_e_ evoked similar effects on cellular excitability, causing severe membrane depolarisation and silencing of firing, presumably through depolarisation blockade (Stojilkovic, 2006; Sankaranarayanan and Simasko, 1996; Kwiecien and Hammond, 1998; Tsaneva-Atanasova et al, 2007). This raises the possibility that, as in hippocampal neurons, NALCN could be responsible for the low [Ca^2+^]_e_-induced depolarisation in the pituitary, providing us with yet another opportunity to functionally identify NALCN in pituitary cells. We test this by assessing the effects of reducing [Ca^2+^]_e_ on pituitary cells excitability.

Consistent with previous studies in neurons, lowering of [Ca^2+^]_e_ (from 2 to 0.1 mM) caused a significant increase in inward leak current in SCR Ctrl pituitary cells (median ± interquartile: −0.8 ± 0.6 pA/pF to −1.5 ± 0.5 pA/pF, n = 15; p < 0.001, Kruskal-Wallis, Figure 6A,C, 6 animals). Subsequent replacement of extracellular Na^+^ with NMDG^+^ reduced the holding current (from −1.5 ± 0.5 pA/pF to −0.3 ± 0.2 pA/pF, n = 15, p < 0.001, Kruskal-Wallis, Figure 6A, C), indicating that the [Ca^2+^]_e_-induced depolarisation is Na^+^ mediated. Lowering [Ca^2+^]_e_ in NALCN KD pituitary cells produced a rise in inward leak current (from −0.37 ± 0.35 pA/pF to −0.5 ± 0.25 pA/pF, n = 13, p < 0.05, Figure 6B, C). However, in contrast, this rise in inward leak current in NALCN KD cells was substantially reduced in comparison with measurements in Ctrl cells (median of Δcurrent-density ± interquartile in SCR Ctrl: −0.6 ± 0.6 pA/pF, n = 13, NALCN KD: −0.13 ± 0.1 pA/pF, n = 13, from 6 mice, p < 0.001, Mann Whitney test, Figure 6D). This indeed indicates that similarly to what is observed in hippocampal neurons, NALCN is involved in pituitary cell sensitivity to reduced extracellular Ca^2+^, and strongly suggest being the mechanism involved in the low Ca^2+^-induced depolarisation observed in pituitary cells.

**Figure 6.**
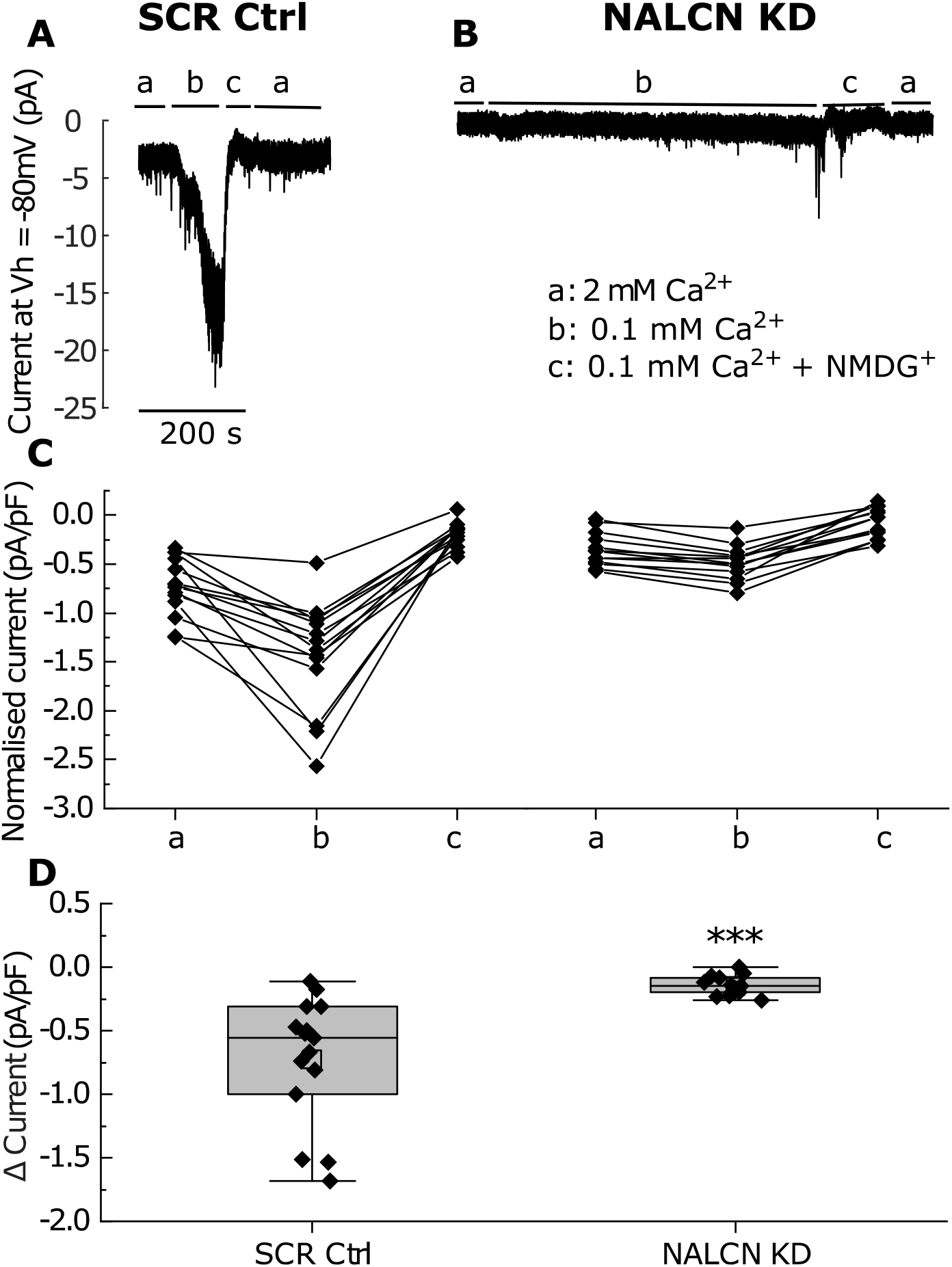
NALCN is sensitive to changes in extracellular Ca^2+^ level. **A-B)** Representative traces of inward leak current in SCR control (SCR Ctrl) and NALCN KD pituitary cells. From baseline conditions (a), reducing extracellular Ca^2+^ level evoked an inward leak current in SCR Ctrl but not in NALCN KD pituitary cells (b). Subsequent replacement of extracellular Na^+^ with NMDG^+^ in reduced extracellular Ca^2+^ returned the inward leak current back to near baseline level (c and a). In **B)** Representative trace of the inward leak current in a NALCN KD pituitary cell; lowering the extracellular Ca^2+^ resulted in a small rise in the inward leak current. Notice that we maintained NALCN KD pituitary cells in low extracellular Ca^2+^ for longer period of time (>200 s) to rule out any further changes in current during application. **C)** Normalised inward leak current in 2mM Ca^2+^ (a), 0.1 mM Ca^2+^ (b), and 0.1 mM Ca^2+^ + NMDG^+^ (c) in SCR Ctrl (left) and NALCN KD (right) cells. **D)** The overall rise in inward leak current after reducing extracellular Ca^2+^ in SCR Ctrl (left) compared to NALCN KD cells (right). ΔMedian ± interquartile in SCR Ctrl: −0.6 ± 0.6 pA/pF, n = 13; NALCN KD: −0.13 ± 0.1 pA/pF, n = 13, from 6 animals, p < 0.001, Mann Whitney test. Box: inter-quartile, Whiskers: range excluding outliers, Central line: median.

## Discussion

We have used a lentiviral-mediated knockdown approach combined with powerful electrophysiology methods and calcium imaging to provide the first compelling evidence that NALCN forms the dominant depolarising conductance in anterior endocrine pituitary cells. Our approach revealed that NALCN channel activity is critical at regulating the electrical and [Ca^2+^]_i_ activity of these cells.

### NALCN regulates the excitability of mouse anterior pituitary cells

Irrespective of cell-types, spontaneous firing in anterior pituitary cells is crucial to maintain normal physiological functions, such as hormone secretion, gene expression and intracellular Ca^2+^ ([Ca^2+^]_i_) oscillations (Kwiecien and Hammond, 1998; Stojilkovic et al, 2010; Fletcher et al, 2018). However, the molecular identity of the major non-selective cationic channels (NSCC) that critically enables spontaneous firing in these cells remains unresolved (Fletcher et al, 2018). Nevertheless, pharmacological studies have revealed that NSCCs are unequivocally Na^+^-dependent and tetrodotoxin (TTX)- resistant/insensitive (Simasko, 1994; Sankaranarayanan and Simasko, 1996; Kucka et al, 2010; Liang et al, 2011; Tomic et al, 2011; Zemkova et al, 2016). Studies using RNA expression profile and non-selective pharmacological manipulations have suggested that transient receptor cation channels (TRPC), as well as hyperpolarisation-activated cyclic nucleotide-gated channel (HCN) may also contribute to the NSCC described in pituitary cells and related cell lines (Tomic et al, 2011; Kucka et al, 2012; Kayano et al, 2019, Kretschmannova et al, 2012). However, without specific receptor blockade and/or genetic interventions (e.g. modification of ion channel expression level) these deductions are largely speculative.

A recent study in the GH3 pituitary cell line has implicated NALCN as a major contributor to the NSCC activity (Impheng et al, 2021). Therefore, we knockdown NALCN activity and used sophisticated electrophysiology and calcium imaging to determine its role in regulating cellular excitability and [Ca^2+^]_i_ oscillations in native anterior endocrine pituitary cells. Remarkably, our results revealed that NALCN channel determines a major Na^+^ leak conductance in these cells, and its activity is critical to sustaining spontaneous firing activity. Our results also show that NALCN is the main contributor to the depolarising NSCC conductance in pituitary cells, without which the cells remained hyperpolarised-silent. In addition, we show that NALCN is crucial for maintaining spontaneous ([Ca^2+^]_i_) oscillations in these cells, and is sensitive to extracellular Ca^2+^ level, as demonstrated in neurons (Lu et al, 2010).

Indeed, we found that NALCN knockdown (KD) pituitary cells are significantly hyperpolarised (by about 15 mV) compared to controls. Moreover, 90% of NALCN KD pituitary cells were completely silent and lost the ability to generate action potential and ([Ca^2+^]_i_) oscillations. Remarkably, restoring the NALCN-like conductance in NALCN KD cells with dynamic clamp rescued cellular resting membrane potential (RMP) and firing activity to normal level. By extension, the removal of such dynamic-clamp mimic of NALCN conductance immediately returned the RMP to hyperpolarised state, silencing the cells.

Together, these results revealed a critical role for NALCN in anterior endocrine pituitary cells, both for maintaining spontaneous firing and ([Ca^2+^]_i_) oscillations.

### A small NALCN conductance sustains a large depolarising drive

The comparison between background Na^+^ inward current in control cells and NALCN KD cells showed that the amount of NALCN conductance knocked down in our experiments was about 0.03 nS. This is consistent with the median of 0.05 nS background Na^+^ conductance required to reactivate silent NALCN KD cells with dynamic clamp. This dynamic clamp-based estimate does not rely on measurements of small noisy currents (a few pA) in voltage-clamp, and therefore provides an estimate that is not affected by poor signal-to-noise ratio (Milescu et al, 2008). The profound effect on electrical activity of such a small conductance is consistent with measurements of high input resistance in pituitary cells, the order of 5 GΩ (Oxford and Dubinsky, 1988), where a very small variation in the current results in a drastic change in RMP. Indeed, with a reversal potential close to 0 mV for this conductance, at a membrane potential of −60 mV, the current through a 0.05 nS NALCN conductance is 3 pA. Therefore, with a membrane resistance of 5 GΩ, the depolarisation due to the current is 15 mV, a value that is remarkably consistent with our finding that NALCN KD cells were hyperpolarised by ~ 15 mV.

We have employed a knockdown approach that is similar to that used in GH3 cells (Impheng et al, 2021), where an estimated NALCN conductance of about 30 pS/pF is reported. This represents a conductance of about three times larger than our estimate in native pituitary cells (10 pS/pF, given our average pituitary cell capacitance of 5 pF). This discrepancy in conductance may be due to the fact that we used a slightly higher extracellular concentration of Mg^2+^ than in Impheng et al, 2021 (1 mM vs 0.8 mM). Given that divalent cations can block NALCN (Chua et al, 2020), this may in part explain the differences seen. A limitation of our work is we have sampled from a heterogenous population of pituitary cells. Thence, different cell-types may express different level of NALCN. If such expression heterogeneity exists, then our estimate of NALCN conductance is an aggregate of the average NALCN conductance expressed by the various cells sampled. Another possibility is that GH3 cells may simply express higher levels of NALCN than primary pituitary cells.

Notably, there is no available estimate for NALCN single-channel conductance in pituitary cells. It is therefore likely that with such knockdown-based approach we are slightly under-estimating NALCN conductance in NALCN KD cells, since there may still be some residual channels expression/conductance. Nevertheless, our estimation of 0.05 nS for NALCN conductance implies a low level of active NALCN channel expression in the the pituitary cell membrane. This may provide support and a parsimonious explanation as to why in our recordings (Figure 4A, 6A), and in the recording of others (Liang et al, 2011) the non-specific cation current is very noisy. Since NALCN has weak voltage sensitivity (Bouasse et al, 2019, Chua et al, 2020) transient blockade by divalent cations, and possibly random channel gating, would result in relatively large current fluctuations if the number of channels is low.

### NALCN activity is modulated by extracellular Ca^2+^ in pituitary cells

Our results also revealed that in mouse anterior pituitary cells, NALCN channels are sensitive to low extracellular Ca^2+^ ([Ca^2+^]_e_), and mediate a low-[Ca^2+^]_e_-induced membrane depolarisation. Consistent with this observation, an increase in [Ca^2+^]_e_ caused hyperpolarisation in cultured rat pituitary somatotrophs through mechanism that does not involved calcium channels (Tsaneva-Atanasova et al, 2007). The [Ca^2+^]_e_-evoked membrane excitation we observed agrees with work in cultured rat lactotrophs and somatotrophs (Sankaranarayanan and Simasko, 1996; Tsaneva-Atanasova et al, 2007), but here we unequivocally revealed that NALCN is directly involved. Our result is also consistent with observation in neurons (Lu et al, 2010; Chu et al, 2003; Bouasse et al, 2019), and work reporting NALCN sensitivity to [Ca^2+^]_e_ blockade in GH3 cells (Impheng et al, 2021).

Two different mechanisms have been proposed to underlie the [Ca^2+^]_e_ sensing capacity of NALCN. First, this may occur through the Ca^2+^-sensing receptor (CaSR) *via* a G_q_-protein dependent pathway that ultimately induces NALCN phosphorylation by protein kinase C (Lu et al, 2010; Lee et al, 2019). Alternatively, NALCN [Ca^2+^]_e_ sensitivity could emerge through direct Ca^2+^ binding within the NALCN pore (Chua et al, 2020). In our experiments, the inward leak current in the majority of the pituitary cells started to increase within 20 seconds after [Ca^2+^]_e_ removal and continued to increase for the next few minutes until it reaches a plateaux (Figure 6A). Thus, although the exact mechanism(s) involved in the NALCN sensing of [Ca^2+^]_e_ in primary pituitary cells is yet to be determined, the timing we observed (with regards to the development of the inward leak current to plateaux level) favours a transduction cascade mechanism over a direct pore effect.

Under both physiological and pathological circumstances, [Ca^2+^]_e_ can drop markedly in the brain and blood serum (Ren, 2011; Ferry et al, 1997). This variation in [Ca^2+^]_e_ is closely linked with adrenocorticotropic hormone (ACTH) secretion from pituitary corticotrophs (Isaac et al, 1984; Fuleihan et al, 1996). In addition, physiological range variations in serum Ca^2+^ is associated with drastic changes in ACTH secretion (Isaac et al, 1984; Fuleihan et al, 1996). Our results, therefore, provide a plausible mechanism for [Ca^2+^]_e_ sensing in pituitary cells, and link this with appropriate hormonal secretion under normal and pathological states.

### NALCN modulation by neurohormones: a likely key player in pituitary cell responses to hypothalamic/feedback signals?

Our finding that NALCN activity has a profound effect on pituitary cell excitability raises the possibility that beyond extracellular Ca^2+^ sensing, NALCN could potentially provide a highly sensitive signalling conduit/target through which hypothalamic neurochemicals/neurohormones could act to regulate hormonal function.

In support, NALCN can be regulated by G-protein dependent and independent signalling pathways (Lu et al, 2009; Swayne et al, 2009; Philippart & Khaliq, 2018), providing a possible intracellular access to its activity by a multitude of receptor systems. Moreover, NALCN is sensitive to hypothalamic neuropeptides, such as Substance P and neurotensin (Lu et al, 2009) which activity have been linked with hormonal regulation (Eckstein et al, 1980). In GH3 cells, NALCN activity promotes basal and TRH-dependent prolactin secretion (Impheng et al, 2021).

These observations are consistent with other *in vitro* work showing that the removal of extracellular Na^+^ to suppress NSCC activity (which our data now identifies as primarily NALCN-driven) in rat pituitary cells negates the ability for growth hormone releasing hormone (GHRH) to evoke growth hormone secretion from these cells (Kato et al, 1988), mostly by preventing (GHRH)-induced Ca^2+^ influx (Lussier et al, 1991; Naumov et al, 1994). This suppression in hormonal secretion is TTX-insensitive and can be rescued by RMP depolarisation with artificially high extracellular K^+^ (Kato et al, 1988). This supports that a Na^+^-dependent depolarising mechanism other than TTX-sensitive voltage-gated Na^+^ channels indeed operates to sensitised hormonal-dependent hormone release by pituitary cells.

Removal of extracellular Na^+^ in murine corticotroph cells also substantially delayed the stimulatory response of these cells to corticotrophin-releasing hormone (CRH) (Liang et al, 2011), presumably by suppressing spontaneous [Ca^2+^]_i_ oscillations in the cells (Tomić et al, 2011). In addition, progesterone and oestrogen have been shown to regulate NALCN expression in human myometrial cells (Amazu et al, 2020), although this remains to be determined in the pituitary. Therefore, altogether, this raises the possibility that hormones and neuropeptides may modulate pituitary NALCN activity in a multifaceted fashion and timescale to influence hormonal release.

In summary, our targeted NALCN knockdown approach, electrophysiology, and calcium imaging methods have highlighted the importance of NALCN activity in supporting appropriate excitability in native pituitary cells. This uncovering may provide an important step in our understanding of pituitary cells excitability and its intersection with hormonal regulation. It may also provide a potential target for therapeutic interventions in endocrine-related disorders that are linked with disruptions in pituitary hormone secretion.

## Materials and Methods

### Primary cell culture

Murine endocrine anterior pituitary cell cultures were prepared between 8:00 and 11:00 AM from wild-type C57BL/6J mice as required. Mice were kept in groups of two to four under standard circumstances at the University of Exeter (UK) animal unit: Lights on at 6:00 AM, lights off at 6:00 PM at 21°C, tap water and food were available *ad libitum*. The adult mice aged between two to six months were selected randomly regardless of their sex. All the animal work was carried out according to the standards of the Home Office (England) and the University of Exeter. Three or four mice were culled via cervical dislocation and then were decapitated in accordance with Schedule 1 procedures. After removing the brain, the pituitary gland was removed from the sella turcica (bony cavity) and placed in a 100×21 mm culture dish (Thermo Fisher Scientific, UK) containing 150 μL 4°C DMEM (Dulbecco’s modified Eagle’s medium with high glucose and 25mM HEPES from Sigma Aldrich (Merck), UK) located on ice. Under a dissection microscope, the intermediate and posterior lobes were removed using a scalpel blade (size 10), and the anterior lobes were chopped to small pieces manually. Subsequently, the chopped tissues were transferred into a 50 mL falcon tube containing 2.5 mL DMEM supplemented with 207 TAME Units/mL trypsin and 36 Kunitz Units/mL DNase I (both from Sigma Aldrich, UK), and then incubated in 37°C water bath for 10 minutes. Every 5 minutes, the tube was swirled to disperse the tissue pieces evenly to achieve a thorough digestion. After 10 minutes, the suspension was gently triturated 20-30 times using a 1 mL pipette tip. At the end of the digestion step, an inhibition solution containing 5 mL DMEM supplemented with 0.25 mg/mL Lima soybean trypsin inhibitor, 100 kallikrein unit aprotinin and 36 Kunitz Units/ml DNase I (Sigma Aldrich, UK) was added to the digestion solution, and the cell suspension was left for a few minutes to inactivate the trypsin enzyme activity. The resulting suspension was finally filtered through a cell strainer with 70 μm nylon mesh (Merck) and was centrifuged at 100×g for 10 minutes. The pellet was resuspended in 500-600 μL DMEM solution and then 60 μL was plated on each 15 mm diameter round coverslip (Thermo Fisher Scientific, UK) in a 12-well plate. After 20 minutes, once the cells were securely attached to the bottom of coverslips, 1 mL of growth medium (DMEM + 2.5% FBS + 0.1 % Fibronectin + 1% antibiotic-penstrep, from Sigma Aldrich, UK) was added to each well and then incubated at 37°C in a 5% CO2 incubator. The culture medium was replaced with antibiotic-free growth-medium 6 hours later. The growth medium was refreshed every two days. 5 μL concentrated suspension of lentivirus was added to the medium in each well of a 12-well plate. Fresh growth medium was substituted 24 hours after transduction. Green fluorescence cells were usually observable 2-3 days after transduction and selected for electrophysiological recording. Each batch of primary pituitary cell culture was utilized for up to 5 days after transduction for electrophysiological recordings and calcium imaging.

### Lentivirus

A microRNA-adapted shRNA based on miR-30 for specific NALCN silencing cloned in the lentiviral pGIPZ plasmid and targeting the 5’-GCAACAGACTGTGGCAATT-3’ region of the rat NALCN-encoding RNA was obtained from a commercial source (Dharmacon #V2LMM_90196). A non-silencing scramble control was used in our experiments (Dharmacon #RHS4346). HpaI/BamHI are the restriction sites in pGIPZ plasmid between which the NALCN silencing shRNA was inserted.

### Electrophysiological recording

Electrophysiological recordings from pituitary cells were performed at room temperature using single-cell amphotericin-perforated patch-clamp technique. The recordings were obtained using an Axopatch 700B amplifier and Clampex 10.1 (Molecular Devices) with a sampling rate of 10 kHz and filtered at 2 kHz (lower pass). Patch pipettes were fabricated from borosilicate glass with filament (outer diameter: 1.50 mm and inner diameter: 0.86 mm, Warner Instrument-multi channel system distributer) and pulled using a micropipette puller (Sutter Instruments, model P-97). Pipette tips were then fire polished and had a resistance ranging from 4 to 6 MΩ. Once a high resistance seal was formed (>10 GΩ), usually within 10 minutes of patching, the access resistance (or series resistance) would reduce to less than 50 MΩ, and then recording started. If the seal resistance was less than 10 GΩ, the cell was discarded. In current clamp mode, series resistance was compensated by Bridge-Balance and was usually less than 40 MΩ. Junction potential was not corrected. In voltage clamp mode, compensated series resistance (60%) was normally less than 50 MΩ and the capacitance of recorded pituitary cells ranged between 4 and 6 pF (electronic compensation was done *via* whole-cell mode of multiclamp 700B). During recording, the cells were constantly perfused using a gravity-driven perfusion system, with a flow rate of 0.5 mL/min with extracellular solution containing (in millimolar) 138 NaCl, 5 KCl, 10 alpha-D-glucose, 25 HEPES, 0.7 Na2HPO4, 1 MgCl2 and 2 CaCl2. The pH was adjusted to 7.4 with NaOH, and the osmolality was 305 mOsmol/L. Patch pipettes were filled with an intracellular solution containing (in millimolar) 10 NaCl, 100 K-Gluconate, 50 KCl, 10 HEPES, and 1 MgCl2. The pH was adjusted to 7.2 with KOH, 295 mOsmol/L. The osmolality of the solutions was maintained by adding an inert ingredient sucrose. 5 μl Amphotericin-B of a stock solution (20 mg/mL in dimethyl sulfoxide) was added to 1 mL of pipette solution to achieve a final concentration of 50 μg/ml. Other concentrations such as 10 μl Amphotericin-B were also tried, however 5 μl resulted in more durability of the recording (all salts and Amphotericin-B were purchased from Sigma Aldrich.

### Dynamic clamp

With dynamic clamp we can effectively change the biophysical property of ion channels (such as activation, inactivation, gating kinetics and conductances) as we choose and investigate how these manipulations affect the pattern of electrical activity in real time. In other words, we can establish the contribution of a type of ion channel to regulating various aspects of electrical activity (Milescu et al, 2008).

In this study, a second computer and an analogue-to-digital acquisition card (DAQ, National Instruments) were installed to run the dynamic clamp module in the software QuB (Milescu et al, 2008). In the current clamp mode of the Axopatch 700B amplifier, the membrane potential (Vm) of a patched cell was recorded in real time and passed to the computer running QuB as an input for a mathematical expression of non-selective cationic leak channels: INs=gNs (V_m_ - E_NS_) which defines the corresponding current (I_NS_) going through them. The sodium leak conductance (g_NS_) was changed manually. The calculated I_NS_ was then injected back to the cell via the same DAQ and then the membrane voltage response was recorded. Thus, the injected NALCN-like current is varied dynamically, unlike conventional current clamp in which the injected current is constant across all time points.

The reversal potential (E_NS_) for this channel was considered zero since this channel is permeable to different monovalent cations (e.g. K^+^ and Na^+^) but primarily to Na^+^ (Lu et al, 2007; Chua et al, 2020).

### Measurement of cytosolic calcium in single pituitary cells

The coverslips with pituitary cells were bathed in the extracellular solution (containing same ingredients as described in the above electrophysiology section) with 2μM fura-2 AM (Thermo Fischer Scientific-# F1221) for 45 minutes at 37°C. The cells were then rinsed three times with the extracellular solution using a 2 mL Pasteur pipette. Following this, the coverslips were mounted onto the recording chamber (volume ~ 0.2 mL) on the stage of an inverted microscope (Nikon eclipse Ti). Cells were constantly perfused with the extracellular solution at room temperature using a gravity-driven perfusion system. Cells were excited every 1 second with alternating 340-nm and 380-nm light beams (20 millisecond exposure time) originating from a Lambda DG-4 wavelength switcher (Sutter Instrument Company). Light intensity was reduced by 50% before hitting the cells using an appropriate filter. The intensity of emitted light was measured at 520 nm and images were acquired by a Hamamatsu digital camera C1344 set to 4 × 4 binning. Hardware control was achieved by TI Workbench software developed by T. Inoue (Tabak et al, 2010). Using this software, regions of interest (ROI) were selected around the cells that were not overlapping with other cells and a single background ROI was selected in an empty space. Pixel values within each region of interest were averaged for both 340 and 380 excitation wavelengths and then subtracted from the background. Following this, a ratio (*r*) was computed according to the formula:

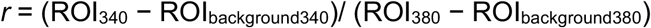

Trace analysis was performed in MATLAB.

### Immunohistochemistry

Pituitary glands were extracted and fixed overnight in 4% PFA in PBS. The following day, 70 μm thick sections were cut using a vibratome (Campden Instruments, UK), placed on poly-L-Lysine coated microscope slides (VWR) and left to dry. Next, the sections were blocked with 10% FBS in PBST (PBS containing 0.01% Triton X-100) for one hour at room temperature. Primary rabbit anti-NALCN antibody at 1:500 dilution (Alomone Labs, Israel, #ASC-022) in 10% FBS/PBST was applied to the sections and incubated in humid chamber at 4 C° degrees overnight. The next day, sections were washed 3 times for 15 min with PBST. Then, the secondary 488-Alexa-Fluor conjugated antibody (at 1:1000, Molecular Probes) was applied in 10% FBS/PBST solution for 1 hour at room temperature, followed by a series of three 15 min washes with PBST. The nucleic acids present in the pituitary gland cells were visualised with TOTO-3 (1:2000, Thermo Fisher Scientific). Sections were incubated with TOTO-3 for 15 min on an agitated surface, then washed three times (each time 10 min) and subsequently mounted with Fluorsave medium (Calbiochem). The controls were the sections where primary antibody was omitted. Images were obtained using a Zeiss LSM 5 Exciter confocal microscope run by Zen software.

### Statistics

We use MATLAB for all statistical analysis and the type of statistical test used in each case is specified in the results.

## Acknowledgements

We would like to thank the members of the University of Exeter Biological Services Unit for their assistance in colony maintenance and husbandry. This work was funded by the University of Exeter PhD studentship to MB, Wellcome Trust ISSF award to JT and JJW, and grant from the Biotechnology and Biological Sciences Research Council (BBSRC) to MDCB (BB/S01764X/1). JJW also acknowledges financial support from the Medical Research Council (MR/N008936/1).

